# An alignment method for nucleic acid sequences against annotated genomes

**DOI:** 10.1101/200394

**Authors:** Koen Deforche

## Abstract

**Motivation:** Biological sequence alignment is fundamental to their further interpretation. Current alignment algorithms typically align either nucleic acid or amino acid sequences. Using only nucleic acid sequence similarity, divergent sequences cannot be aligned reliably because of the limited alphabet and genetic saturation. To align divergent coding nucleic acid sequences, one can align using the translated amino acid sequences. This requires the detection of the correct open reading frame, is prone to eventual frame shift errors, and typically requires the treatment of genes separately. It was our motivation to design a nucleic acid sequence alignment algorithm to align a nucleic acid sequence against a (reference) genome sequence, that works equally well for similar and divergent sequences, and produces an optimal alignment considering simultaneously the alignment of all annotated coding sequences.

**Results:** We define a *genome alignment score* for evaluating the quality of an alignment of a nucleic acid query sequence against a reference genome sequence, for which coding sequence features have been annotated (for example in a GenBank record). The genome alignment score combines the a ne gap score for the nucleic acid sequence with an a ne gap score for all amino acid alignments resulting from coding sequences in open reading frames contained within the query sequence. We present a Dynamic Programming algorithm to compute the optimal global or local alignment using this genomic alignment score and provide a formal proof of correctness. This algorithm allows the alignment of nucleic acid sequences from closely related and highly divergent sequences within the same software and using the same parameters, automatically correcting any eventual frame shift errors and produces at the same time the aligned translated amino acid sequences of all relevant coding sequence features.

**Availability:** The software is available as a web application at http://www.genomedetective.com/app/aga and as command-line application at https://github.com/emweb/aga

## 1 Introduction

Obtaining nucleic acid sequences that span partial or full genomes are becoming cost-effective with recent advances in sequencing technology. This enables new applications that use use this genomic sequence data from known and unknown species in clinical or environmental samples [7] [15].

Alignment, as part of a database similarity search, or against a specific reference genome, is typically a first step in the interpretation of these sequences. Current alignment methods however work either on the nucleic acid or amino acid sequence, but typically do not consider both simultaneously. The benefit of aligning amino acid sequences is that they are usable for more diverged sequences, but they require the correct detection of open reading frames and a priori correction of eventual frame shift errors that interfere with proper translation. By considering only amino acid sequence similarity, the more sensitive evolutionary information within synonymous substitutions is however lost.

We describe here an algorithm (AGA) which computes the optimal pairwise alignment of an (unknown) nucleic acid sequence against a reference sequence, using a score that combines nucleic acid similarity and amino acid similarity. Amino acid similarity is based on translated Coding DNA Sequence (CDS) annotations of the reference sequence. In a typical application, the reference sequence would be a complete genome sequence annotated with the location of coding sequences of contained proteins or polyproteins (see for example Figure 2). The proposed method can deal with multiple coding sequences in different open reading frames, which use either the forward or the reverse complemented strand, and which may be composed of different regions that are spliced together. Open reading frames may also verlap, which is not uncommon for compact viral genomes.

The alignment of coding nucleic acid sequences using amino acid sequence similarity has been considered before [5] [1] [18] [18] [2] [19] [11] [13] [8] [17]. The method outlined in this paper di ers with this previous work in the sense that it considers specifically the problem of how to optimally align a nucleic acid sequence against an annotated (reference) genome, considering simultaneously nucleotide simularity and amino acid sequence simularity in the annotated coding sequences. The method results in alignments with a minimum number of frame shifts in coding sequences and with gaps preferably at codon boundaries, by penalizing both such events in the scoring function that it optimizes. By combining similarity of the nucleic acid sequences and similarity of coded amino acid sequences, the method can optimally align sequences to a reference genome, regardless of whether they are highly similar or distant to the reference genome.

Applications such as phylogenic tree reconstruction, sequence similarity evaluation, read or contig mapping towards a reference sequence, or determination of nucleic acid and amino acid substitutions for geno-typic/phenotypic assocations, all depend on a high quality alignment. We believe that AGA is useful to most of these applications.

## 2 Approach

We define a *Genome Alignment Score* which scores the quality of a local or global alignment of a nucleic acid *Query* sequence against a *Reference Genome*. In this context, a reference genome is a nucleic acid sequence annotated with *Coding DNA Sequence* information. Each coding sequence indicates the location of one or regions in the genome that jointly translate to a protein or polyprotein. The coding sequence may be based on a single region or may span multiple regions that are spliced together. Each region is part of one of three forward or reverse complemented open reading frames within the genome sequence. Coding Sequences may also overlap, sharing the same or different open reading frames, which is not unusual for compact viral genomes.

The genome alignment score is based on (a) the alignment score of the nucleic acid sequence using a traditional nucleic acid substitition matrix score, with affine gap open and gap extension penalties [16], (b) the alignment score of each of the covered coding sequences using a traditional amino acid substition matrix score, with affine gap open an gap extension penalties, and (c) additional penalties for frameshift insertions and deletions, and for insertions and deletions that do not align with codon boundaries.

By defining the genome alignment score in this way, it can be used to compare the similarity of a nucleic acid sequence to multiple reference genomes. By including the alignment score of the nucleic acid sequence itself, the genome alignment score is applicable to sequences that are highly similar to the reference, having for example only meaningful differences in their nucleic acid sequences, but virtually no changes in their amino acid sequences. By including the alignment score of all covered coding sequences as well, the alignment score is in particular also applicable to highly divergent sequences, which may have lost much of their similarity at the nucleic acid sequence, but which still share some similarity in their protein sequences, especially when considering all coding sequences (covered by the query sequence) simultaneously. Finally, by considering the possibility of (likely erroneous) insertions or deletions that cause frame shifts, the alignment score is suitable to align sequences obtained from sequencing techniques that are prone to such sequencing errors without misusing frameshift mutations artificially as a means to optimize amino acid sequence similarity.

We show how a Dynamic Programming algorithm can be designed which computes the optimal local or global alignment subject to maximizing the genome alignment score. This work thus builds further on the optimal alignment algorithms first proposed by Needleman-Wunsch [16], Smith-Waterman [16], and Gotoh [4], by expanding the induction state with additional state parameters.

## 3 Methods

### 3.1 Notation

Let the two nucleic acid sequences be a reference genome G = g_1_g_2_…g_*M*_ and a query sequence B = b_1_b_2_…b_*N*_. For the reference genome G, information on coding sequences are available, which are for example the CDS annotations from a GenBank record. This information can be represented as a list of codons in which the nucleotide at position *m* participates: *C_m_* = {[*c*_*m*,1_, *r*_*m*,1_], [*c*_*m*,2_, *r_m_*,2], …, [*c_m,t_, r_m,t_*]} where 1 ≤ *c*_*m,i*_ ≤ 3, indicating the position of the nucleotide in a codon, and *r_m,i_* a boolean indicating whether the codon is in the forward strand or in the reverse complementary strand. A splice site does not necessarily occur at a codon boundary, and in that case the nucleic acids that are translated to the codon may be scattered through the genome. To allow the algorithm below to process the sequences sequentially from start to end, we exclude such codons from the scoring models. We define *t*_*m*_ = |*C*_*m*_|, the number of codons in which the nucleotide takes part. A value of *t*_*m*_ > 1 indicates that there are multiple overlapping coding sequences at the given nucleotide position *m*, possibly in different open reading frames. We denote as *T_i_*(a, *r*) the translation of the codon a_*i*_a_*i*+1_a_*i*+2_ to an amino acid in the forward or reverse complementary strand depending on the value of *r*.

### 3.2 Genome alignment score

For a nucleic acid sequence alignment *A*_na_(G, B), we define a *genome alignment score S*_ga_{*A*_na_(G, B)} that is based on a nucleic acid sequence alignment score *S*_na_{*A*_na_(G, B)} for the nucleic acid sequence alignment *A*_na_(G, B) itself, and the amino acid sequence alignment score *S*_aa_{*A*_aa_(X, Y)} for each amino acid sequence alignments *A*_aa_(X, Y) that results from translation of the aligned sequences *G* and *B* according to the coding sequence annotations of *G*.

The alignment scores *S*_na_ and *S*_aa_ are of the same form but use a distinct set of parameters. They score a match in the alignment according to a substitution weight matrix *W*, and a gap of length *k* in the query or reference sequence using an affine gap model: *w*_*k*_ = *p*_*u*_(*k* − 1) + *p*_*v*_ with *p*_*v*_ ≤ 0 the penalty for opening a gap, and *p*_*u*_ ≤ 0 is the cost for extending a gap. The incremental cost for a gap is then

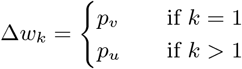

Because the genome alignment score, and the alignment algorithm described below, consider the insertion of gaps anywhere in the nucleic acid sequence, and gaps of any length, we need to define how these gaps within a coding sequence translate to gaps in the amino acid sequence. We make the choice to consider a gap of length *k* with 3(*n* − 1) < *k* ≤ 3*n* as a gap of length *n* in the amino acid sequence, regardless of whether the gap aligns with a codon boundary. For practical reasons, one typically prefers gaps to align with codon boundaries because otherwise they render the interrupted start and end codons as non-translatable, which lack a proper notation and scoring at the amino acid sequence level (usually denoted as an ‘X’ in the resulting sequence alignment). Therefore a misalignment penalty is added whenever a gap starts within a codon, disrupting proper translation of that codon. More formally, a penalty *p_m_* ≤ 0 is added whenever a gap opens after position g_*m*_, either in the query or reference sequence, when *C_m_* contains a codon position *c*_*m,i*_ ≠ 3.

The amino acid sequence alignment score *S_AA_* by itself does does not consider insertions and deletions in the underlying nucleic acid sequence that do not occur in a multiple of three. These however cause frameshift mutations which change the translation profoundly and thus has a large impact on the amino acid sequence alignment. Frame shifts are unexpected in a viable coding sequence, but they are not uncommon as a consequence of sequencing errors and thus their possibility needs to be considered (with low probability) to obtain a better alignment. The introduction of a frame shift needs thus to be weighted against the quality of the amino acid sequence alignment. Thereofre, in the genome alignment score, a frame shift penalty *p_f_* ≤ 0 is added for a gap of length *k* with (*k* mod 3) ≠ 0.

Finally, the genome alignment score S_ga_ is defined as a weighted sum of the nucleic acid sequence alignment score and the amino acid sequence alignment scores, using a weight *w*_aa_ for the amino acid alignment score contribution.

### 3.3 Algorithm

We now describe an algorithm that calculates the optimal alignment, maximizing the above genome alignment score, using Dynamic Programming. The algorithm extends the idea developed in [4] to expand the induction state with additional state matrices to properly accomodate the affine gap cost in the induction step. In particular, in [4] the induction state was defined as [*D_m,n_*, *P_m,n_*, *Q_m,n_*] in which *D_m,n_* is the best score for an alignment of g_1_g_2_…g_*m*_ versus b_1_b_2_…b_*n*_, *P_m,n_* the best score for an alignment of the same sequences but ending with a gap after g_*m*_, and *Q_m,n_* the best score for an alignment of the same sequences but ending with a gap after *b_m_*. These additional two matrices *P_m,n_* and *Q_m,n_* allow the algorithm to properly evaluate the score for a gap taking into account the different cost for a gap open and gap extension. With the genome alignment score as defined above, the cost for opening a gap will be different from a gap extension, not only because of the affine gap penalty present in the nucleic acid sequence and amino acid sequence alignment scores, but also because opening a gap may break one or more codons and cause a codon misalignment cost *p_m_*. But also the cost of a gap extension differs from one to another since the length of the gap influences the occurence of a frame shift penalty *p_f_*, and depending on the position relative to a codon, a gap penalty may be added to the amino acid alignment.

We define as induction state 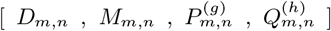 with 1 ≤ *g* ≤ 3 and 1 ≤ *h* ≤ 3 where for an alignment of g_1_g_2_…g_*m*_ versus b_1_b_2_…b_*n*_:

- *D_m,n_* is the best score overall;
- *M_m,n_* is the best score for an alignment ending with a match;
- 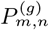 is the best score for an alignment ending with a gap of length *k* = 3*i* + *g* after g_*m*_ with *i* ≥ 0 in the reference sequence;
- 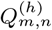 is the best score for an alignment ending with a gap of length *k* = 3*j* + *h* after b_*n*_ with *j* ≥ 0 in the query sequence.

At each induction step *m, n* the induction state is updated using an incremental computation of the genomic alignment score. Let Δ*d_m,n_* be the incremental score for extending the alignment of g_1_g_2_…g_*m*−1_ with b_1_b_2_…b_*n*−1_ with a match. Δ*p_m,n_*(*k*) is the incremental score for extending an alignment of g_1_g_2_…g_*m*_ with b_1_b_2_…b_*n*−1_ by opening a gap (*k* = 1) or extending a gap in the reference sequence to length *k*. Likewise let Δ*q_m,n_*(*k*) be the incremental score for extending an alignment of g_1_g_2_…g_*m*−1_ with b_1_b_2_…b_*n*_) by opening a gap (*k* = 1) or extending the gap in the query to length *k*.

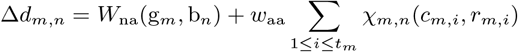

The sum component in the above equation adds the amino acid substitution score for each codon which starts at position *m*, and thus for which *c_m,i_* = 1.

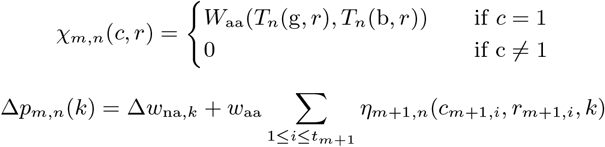

The sum component adds a score component *η* for each amino acid sequence which may be affected by the gap.

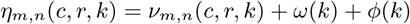

The codon breakage cost *ν* undoes an amino acid substitution weight previously added at the beginning of the codon, plus adds a misalignment penalty:

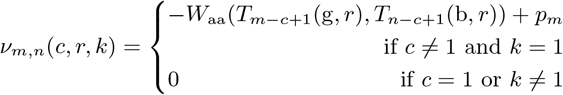

An incremental amino acid open gap cost is added when opening the gap, and an amino acid gap extension cost is added for every additional 3 nucleic acid gaps, according to the affine gap model:

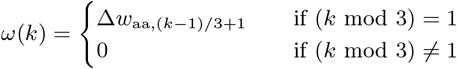

A frame shift penalty is incrementally updated based on the length of the gap *k*:

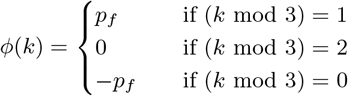

Likewise we can define the incremental score for opening or extending a gap in the query sequence:

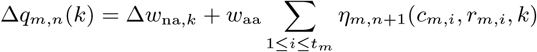

At each induction step, we use the above incremental score update functions to update the induction state (Figure 1):

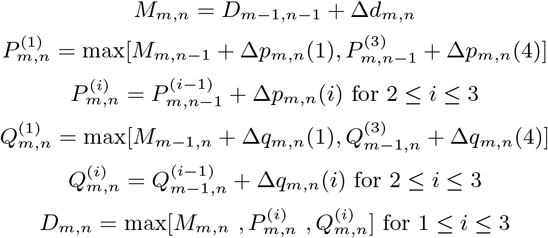

**Figure 1:**
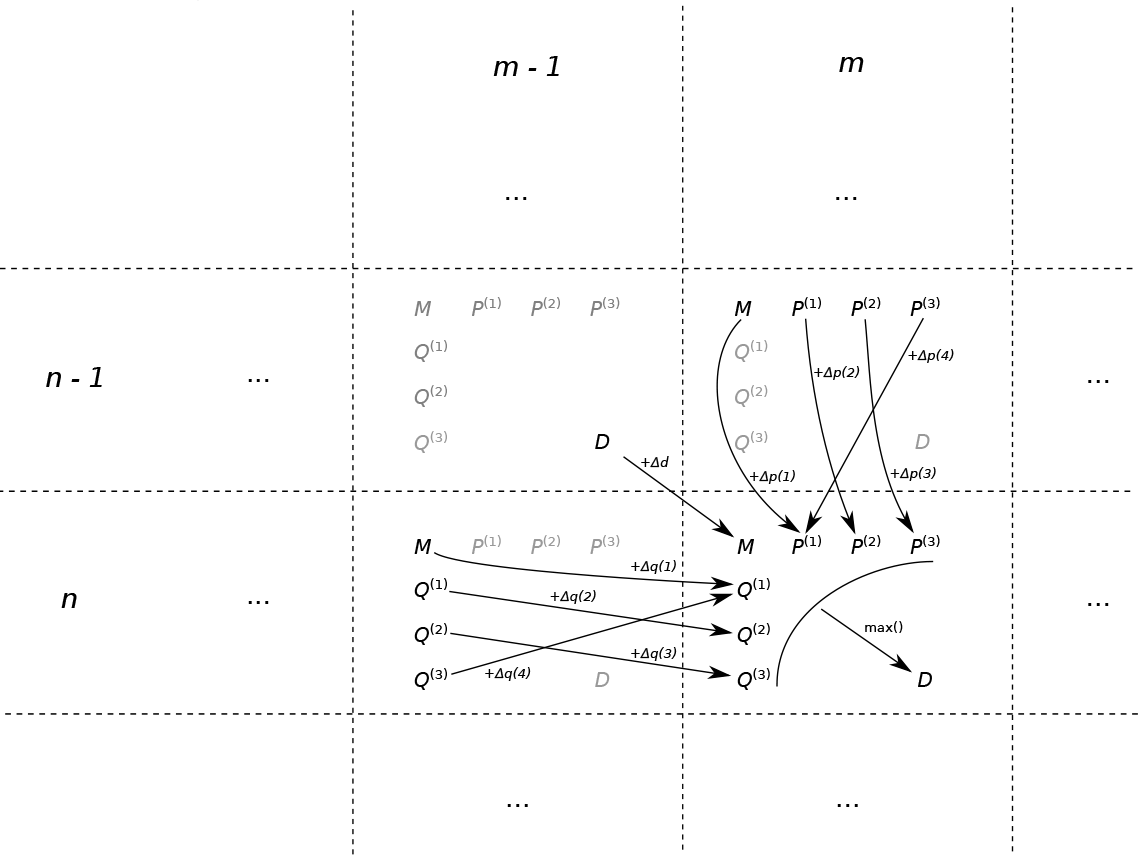
Data dependencies in induction rule

The optimal alignment is retrieved by tracking back the path through the induction state matrix from *D_m,n_* to *D*_0,0_ following back the path that led to the optimal solution.

### 3.4 Proof

At each induction step *m, n* we need to prove that the definitions of the induction state parameters are satisfied.

The definition of *M_m,n_* is satisfied since Δ*d_m,n_* only depends on *m* and *n* and thus the optimal alignment ending in a match is the alignment that extends the optimal alignment of g_1_g_2_…g_*m*−1_ versus b_1_b_2_…b_*n*−1_ with score *D*_*m*−1,*n*−1_.

The defintions of 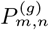 and 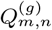 for *g* = 2 or *g* = 3 are satisfied because the incremental gap cost terms Δ*p_m,n_*(*k*) and Δ*q_m,n_*(*k*) depend only on *m*, *n*, and gap length *k*. When extending a gap (*k* ≠ 1), it can easily be verified that the following equalities exist:

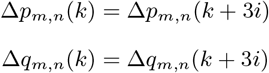

This means that the cost for extending a gap to length *k* = 3*i* + *g* is the same for any value of *i*, and thus the highest score for an alignment ending with a gap of length *k* = 3*i*+*g* is the highest score for an alignment ending with a gap of length *k* = 3*i* + (*g* − 1), incremented with the gap extension cost for *k*.

The definitions of 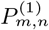 and 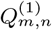 are satisfied because the algorithm uses the maximum score considering either the opening of a gap or extending of a gap of length *k* = 3*i*. The cost for opening a gap Δ*p_m,n_*(1) and Δ*q_m,n_*(1) depend only on *m* and *n*, and thus the highest score for ending the alignment with a gap of length *k* = 3*i* + 1 is either the cost for opening a gap added to the highest score for ending with a match, or for extending a gap of length *k* = 3*i*.

Finally, the optimal alignment score *D_m,n_* is defined as the maximum alignment score considering the different options of ending the alignment in a match, or ending with a gap in the reference of all possible lengths, or ending with a gap in the query of all possible lengths.

## 4 Implementation

The above algorithm was implemented in C++11 as a standalone commandline tool (AGA). The inputs are a reference genome (GenBank record) and a query nucleic acid sequence (FASTA file). The implementation does not keep the entire induction state matrix in memory by encoding the backtrace information within the induction state variables, and is thus in practice suitable for genome lengths up to 10^6^ base pairs (most viral genomes), provided sufficient computation time.

Through command-line parameters, the following parameters of the algorithm may be configured: the choice of nucleic acid and amino acid substitution weight matrices *W*_na_ and *W*_aa_, the affine gap parameters *p*_*u*,na_, *p*_*v*,na_, *p*_*u*,aa_, *p*_*v*,aa_, parameters to weight the nucleic acid versus amino acid score *w*_*aa*_, and frame shift penalty *pf* and misalignment penalty *p*_*m*_. For the nucleic acid substition weight matrix, a score for a match and a score for a mismatch can be configured. As amino acid substitution weight matrices the tool offers the choice between BLOSUM30 and BLOSUM62 [6]. AGA can compute an optimal local or global alignment.

The tool outputs the nucleic acid sequence alignment and all amino acid sequence alignments of coding sequences (after eventual frame shift corrections). For each alignment, it outputs the corresponding score and provides various statistics (coverage length, number of matches, identities, indels, frame shifts, and codon misalignments).

## 5 Evaluation

To assess the benefit of the algorithm, we compared it to an implementation of a an optimal nucleic acid sequence alignment which doesn’t take into account CDS features (EMBOSS needle [14]), and other alignment algorithms designed to take into account CDS features (Table 1). For the comparison, we evaluated how these algorithms perform to align different HIV and other primate lentivirus genome sequences against the HIV-1 reference sequence (Table 2).

**Table 1:** Algorithms whose performance was compared to AGA.

| Name | Version | Score | Input | Description |
| --- | --- | --- | --- | --- |
| needle | 6.6.0 | NT | pair-wise | no special support for CDS |
| MACSE | 0.9b1 | AA | multiple | one CDS, minimizes frame shifts |
| codaln | 1.0 | NT + AA | multiple | multiple CDS |

**Table 2:** Sequences used in the evaluation of AGA.

| Name | Accession | Length | Description |
| --- | --- | --- | --- |
| HXB2 | NC_001802 | 9181 | HIV-1 reference strain (Subtype B) |
| hiv1b | AY835761 | 9824 | HIV-1 Subtype B complete genome |
| hiv1c | U46016.1 | 9031 | HIV-1 Subtype C complete genome |
| siv | KF304708.1 | 9449 | SIV complete genome |
| hiv2 | KP890355.1 | 9480 | HIV-2 complete genome |

HIV-1 was chosen for this evaluation since it has a complex genome organization (Figure 2), which is not uncommon for RNA viruses, with multiple overlapping reading frames, some of which are joined by different Coding DNA Sequence (CDS) annotations of HIV-1, as derived from the GenBank record NC 001802 (HIV-1 reference sequence) genomic segments, and including one that uses the complementary strand (for the *asp* gene). Although one routinely will align HIV-1 sequences to the reference genome, also the alignment of HIV-2 and SIV strains against HIV-1 may be useful to unravel genetic causes for the different phenotypic properties of these viruses [12] [3].

**Figure 2:**
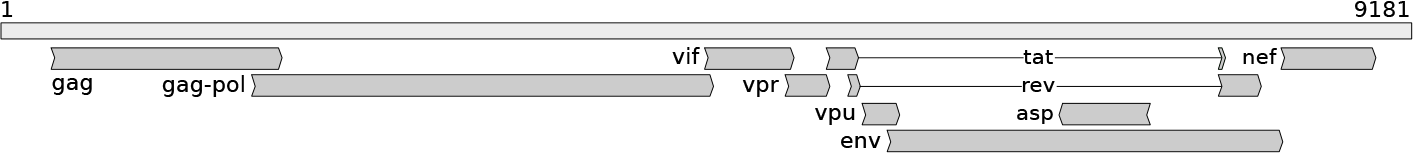
HIV-1 Genome Organization

To evaluate the quality of each alignment, we used as meaningful statistics number of frame shifts introduced within coding sequences, the number of amino acids not aligned at a codon boundary, and the affine gap model score for the nucleic acid sequences and all amino acid sequences (excluding frame shift and misalignment penalties). The run time performance was also compared on a Dell XPS laptop.

Tables 3 and 4 detail value parameters used for running the different algorithms in the comparison, which we tried to make comparable despite differences in scoring models.

**Table 3:**
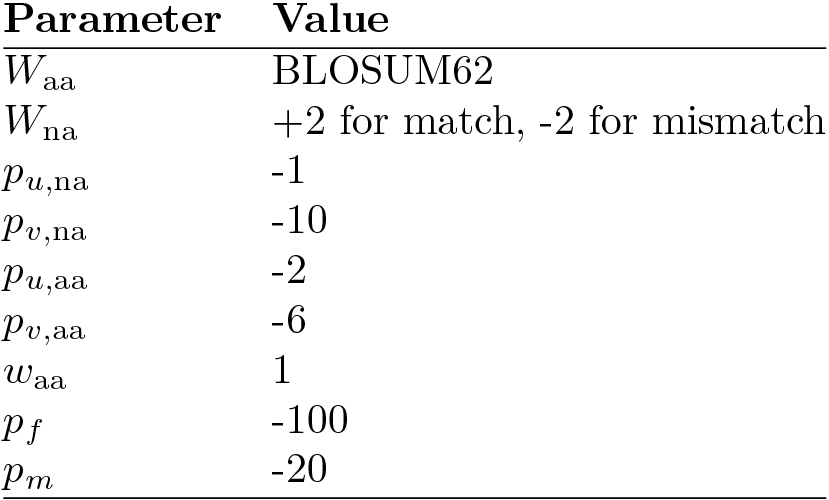
Parameter values used for AGA in its evaluation.

**Table 4:**
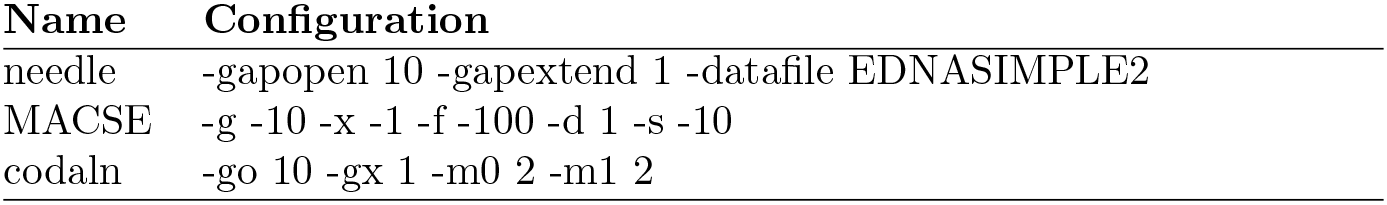
Command-line arguments used for other algorithms used in the evaluation.

| Name | Configuration |
| --- | --- |
| needle | -gapopen 10 -gapextend 1 -datafile EDNASIMPLE2 |
| MACSE | -g -10 -x -1 -f -100 -d 1 -s -10 |
| codaln | -go 10 -gx 1 -m0 2 -m1 2 |

We also compared the different alignments using an alignment viewer that shows simultaneously the nucleic acid sequence alignment and all coding sequence alignments (this alignment viewer is also part of the web version othe tool).

## 6 Results

AGA produced alignments with high nucleic acid and amino acid alignment scores while introducing only a minimum amount of frame shifts and codon misalignments within coding sequences (Table 5). Because MACSE makes the assumption of a single CDS, it introduces a frame shift near the beginning or the end of a coding sequence region, or within an overlapping region, to jump to the new gene open reading frame, even for the alignment of the highly similar HIV-1 subtype B sequence, but otherwise also effectively minimizes the number of frame shifts. Codaln on the other hand, like AGA, uses its knowledge of open reading frames to minimize frame shifts within Coding Sequences, but the alignments are of a lower quality (at both the nucleic acid and amino acid level) because of an inaccurate estimation of open reading frames in the query sequence.

**Table 5:** Alignment quality of AGA compared to other algorithms

| Sequence | Algorithm | # FS | # MA | NT Score | AA Score |
| --- | --- | --- | --- | --- | --- |
| hiv1b | AGA | 0 | 10 | 16182 | 18130 |
|  | needle | 3 | 20 | 16219 | 18054 |
|  | MACSE | 13 | 29 | 15521 | 18013 |
|  | codaln | 3 | 16 | 15615 | 18035 |
| hiv1c | AGA | 1 | 19 | 12478 | 16127 |
|  | needle | 22 | 54 | 12525 | 16097 |
|  | MACSE | 13 | 42 | 11772 | 15813 |
|  | codaln | 4 | 21 | 11930 | 15907 |
| siv | AGA | 2 | 23 | 1497 | 8536 |
|  | needle | 146 | 298 | 1980 | 8159 |
|  | MACSE | 13 | 66 | 62 | 8301 |
|  | codaln | 5 | 38 | 34 | 7552 |
| hiv2 | AGA | 1 | 30 | 1164 | 8080 |
|  | needle | 161 | 338 | 1650 | 7472 |
|  | MACSE | 18 | 78 | 213 | 7799 |
|  | codaln | 1 | 27 | -85 | 6983 |

Performance of other algorithms compared to AGA for aligning four genome sequences against the HIV-1 reference strain (HXB2): # *FS* is the number of frame shifts caused by indels within Coding DNA Sequences (CDS) in either the reference or query sequence; # *MA* is the number of indels not aligned to a codon boundary; *NT Score* is the nucleic acid affine gap model score of the alignment; *AA Score* is the sum of all amino acid affine gap model scores of the translated CDS alignments, after inserting additional gaps to correct eventual frame shift errors.

EMBOSS needle produces alignments with a maximum nucleic acid alignment score, as can be expected, but it will introduce frame shifts and codon misaligned gaps in order to optimize its nucleic acid sequence alignment.

AGA as well as the other algorithms tested, implement a Dynamic Programming solutions with a time complexity *O*(*MN*), and with the exception of MACSE, all had a similar run time performance (Table 6). Taking into account the possibility of multiple overlapping open reading frames, AGA has an *O*(*MNT*) time complexity, where *T* is the maximum amount of overlapping coding sequences within the genome: *T* = max_*i*_ *t*_*i*_.

**Table 6:** Runtime performance of AGA compared to other algorithms

| **Algorithm** | **Run time (s)** |
| --- | --- |
| AGA | 6.3 |
| needle | 4.5 |
| MACSE | 185.2 |
| codaln | 5.5 |

Run time performance of other algorithms compared to AGA for aligning an HIV-1 Subtype B full genome sequence (hiv1b, 9824 bps) against the HIV-1 reference strain (HXB2, 9181bps) on a Dell XPS (Intel Core i7-7500U CPU @ 2.70GHz).

## 7 Discussion

The proposed genomic alignment score combines the affine gap model score of the nucleic acid sequence alignment with all affine gap model scores of amino acid alignments. These scores are however not independent: a gap in an amino acid sequence alignment will also correspond to a gap in the nucleic acid sequence alignment. A matching amino acid in the amino acid sequence alignment may correspond to matching codons in the nucleic acid sequence alignment. Nevertheless the approach to allow gaps to be penalized at both levels was chosen since the cost for the introduction of the gaps is also weighted against the score for character matches in the alignments at each level. The parameter *w*_aa_ can be used to give more or less weight to the amino acid sequence alignments compared to the nucleic acid sequence alignment and a suitable value will depend also on the nature of both scoring models since these are dimensionless numbers that are not necessarily of the same order of magnitude and thus comparable.

The original motivation for affine gap costs in nucleic acid sequence alignments was in part to avoid gaps that introduce frame shifts [16]. Since in the genomic score the amino acid sequence alignment score is included, it may have become redundant. AGA can also implement a linear gap model by setting *p*_*u*,na_ = *p*_*v*,na_.

From the results (Table 5), it can be seen that AGA can still generate indels that are not aligned with a codon boundary in one of the coding sequences. Provided a sufficiently high penalty for this, this will only happen in regions of overlapping reading frames where the gap can only be codon-aligned with one of the amino acid sequences.

A number of algorithms have been proposed which align coding nucleic acid sequences by back-translation of the corresponding amino acid sequence alignment [1] [18] [19] [18] [2]. Such methods however are limited to coding sequences for a single protein or polyprotein (and cannot deal with overlapping open reading frames), cannot easily deal with frame shift errors that prevent the translation in the first place, and disregard the contribution of nucleotide similarity entirely.

To explicitly consider the possibility of frame shifts, another class of algorithms more similar to AGA have been proposed, which therefore modify nucleic acid sequence alignment algorithms to take into account the translation, but still allow for frame shifts [13], [5], [17]. Like AGA, these algorithms use a Dynamic Programming induction matrix to compute an optimal alignment subject to a scoring function, and different results are caused by different assumptions embedded in their scoring functions.

Codaln [17] implements an algorithm which, like AGA, can read the annotations of coding sequences from a GenBank record. To align an unknown query sequence against an annotated genome, it will first search for open reading frames in the query sequence, which are used in a second step in the scoring function of a Dynamic Programming alignment algorithm. In our results (Table 5), we found that the lower alignment scores, and erroneous frame shifts, of its alignments were caused by errors in these estimated open reading frames.

MACSE [13] penalizes frame shifts but assumes that the entire sequence is a single Coding Sequence which needs to be translated with a minimum of frame shifts and stop codons into a (pseudo-)protein, while optimizing the resulting amino acid alignments. Although we included MACSE in our evaluation, it is thus not well suited to align entire genome with multiple, partially overlapping, open reading frames, some of which may be using the complementary strand.

Future work could be to extend the scoring model to multiple sequence alignment, considering then that one of the included sequences is a reference genome with CDS annotations. This could use the progressive combination of pairwise alignments as originally implemented in CLUSTALW [10], or other heuristic approaches such as implemented in MAFFT [9] or MUSCLE [9].

## 8 Conclusion

We presented an optimal solution to a fundamental problem in biological sequence analysis, namely how to best align an unknown nucleic acid sequence against a (reference) genome, considering simultaneously similarity at the nucleic acid and amino acid sequence level, and condidering possible frame shifts and gaps causing codon misalignment, but scoring such events with a user-defined penalty.

The proposed method is generally applicable to align any nucleic acid sequence against any genome for which Coding Sequence feature annotations are available. In practice, it is especially well suited to RNA virus sequences, since they are generally rapidly evolving and also typcally have compact viral genomes. By considering amino acid sequence similarity across all Coding Sequences, the method can overcome the large diversity caused by the high rate of evolution, and deals properly with overlapping reading frames common to these viruses.

The method has been implemented in a software package AGA which is available as a command-line package or can be used through a simple web page.

## Funding

The VIROGENESIS project receives funding from the European Unions Horizon 2020 research and innovation programme under grant agreement No 634650.

